# A Robust Targeted Sequencing Approach for Low Input and Variable Quality DNA from Clinical Samples

**DOI:** 10.1101/123117

**Authors:** Austin P. So, Anna Vilborg, Yosr Bouhlal, Ryan T. Koehler, Susan M. Grimes, Yannick Pouliot, Daniel Mendoza, Janet Ziegle, Jason Stein, Federico Goodsaid, Michael Y. Lucero, Francisco M. De La Vega, Hanlee P. Ji

**Author notes:** These authors contributed equally to this work (Austin P. So, Anna Vilborg). Corresponding authors: **Correspondences:** Anna Vilborg, Hanlee P. Ji.

## Abstract

Next-generation deep sequencing of gene panels is being adopted as a diagnostic test to identify actionable mutations in cancer patient samples. However, clinical samples, such as formalin-fixed, paraffin-embedded specimens, frequently provide low quantities of degraded, poor quality DNA. To overcome these issues, many sequencing assays rely on extensive PCR amplification leading to an accumulation of bias and artifacts. Thus, there is a need for a targeted sequencing assay that performs well with DNA of low quality and quantity without relying on extensive PCR amplification. We evaluate the performance of a targeted sequencing assay based on Oligonucleotide Selective Sequencing, which permits the enrichment of genes and regions of interest and the identification of sequence variants from low amounts of damaged DNA. This assay utilizes a repair process adapted to clinical FFPE samples, followed by adaptor ligation to single stranded DNA and a primer-based capture technique. Our approach generates sequence libraries of high fidelity with reduced reliance on extensive PCR amplification - this facilitates the accurate assessment of copy number alterations in addition to delivering accurate SNV and indel detection. We apply this method to capture and sequence the exons of a panel of 130 cancer-related genes, from which we obtain high read coverage uniformity across the targeted regions at starting input DNA amounts as low as 10 ng per sample. We further demonstrate the performance of this assay using a series of reference DNA samples, and by identifying sequence variants in DNA from matched clinical samples originating from different tissue types.

## INTRODUCTION

Next-generation sequencing **(NGS)** with targeted gene panels has seen general adoption as a diagnostic and screening tool for a wide variety of disorders.^1^ Clinical applications include: 1) identifying germline variants, such as single nucleotide polymorphisms **(SNPs)** and structural variants **(SV)** related to hereditary disorders, and 2) identifying somatic mutations and other genetic aberrations in cancer that may have implications for treatment and prognosis.^2^ Cancer somatic mutations frequently occur at low variant allelic fractions **(VAF)** and are more difficult to detect from biopsy samples.^3^ The use of targeted gene panels has multiple advantages in all of these cases. Deep sequencing of genes and other clinically-actionable genomic targets results in higher read coverage, oftentimes in the thousand-fold range, and as a result, improves the confidence and the analytical detection limit of variant alleles.^4^ This is particularly valuable for analyzing clinical samples that are composed of genetic mixtures, such as solid tumors where multiple clones of cancerous cells are mixed with normal stromal components.

A major challenge for diagnostic sequencing is the variable quality of the genomic DNA obtained from clinical samples. This variability arises in part from the adverse effects of processing applied to samples upon the integrity of DNA.^5^ Specifically, the vast majority of clinical tumor biopsies undergo formalin fixation and paraffin embedding **(FFPE)** to facilitate histopathologic examination. Unfortunately, this archival process modifies nucleotides, generates chemical crosslinks, and can lead to degradation of the DNA over time. Consequently, DNA purified from FFPE is often fragmented and contains a significant proportion of damaged and single stranded molecules.^6^ As a result, molecular diagnostics based on FFPE DNA often require a high degree of optimization, and assay failures are significantly more frequent than instances where higher quality DNA is available.^5^ Indeed, many methods now employ DNA quality control criteria to reject samples to mitigate test failures due to sample quality.^7^ While increasing the success of the diagnostic assay, these exclusion criteria eliminate some samples that may be of clinical significance and value.

To address the challenges of efficiently detecting variants of low VAF from FFPE material, we developed a targeted sequencing approach termed Oligonucleotide-Selective Sequencing **(OS-Seq).** This approach has multiple features that facilitate its application to diagnostic targeted sequencing of DNA from a variety of clinical samples.^8,9^ In particular, we have developed an in-solution version of OS-Seq that provides a streamlined and efficient process for targeted sequencing without the need for flow cell modification, as required in the original version. This assay was optimized for the sequencing of clinical samples of variable quality, and draws upon methods for sequencing ancient DNA samples. ^10,11^ In particular, in-solution OS-Seq involves a pre-processing step that excises damaged bases without corrective repair, followed by a highly efficient single stranded adapter ligation process. The efficient ligation allows for the conversion of all nucleic acid species – regardless of quality and quantity – into partial sequencing libraries with a single adapter. Following this ligation process, target-specific multiplexed primer annealing and extension of the genomic targets on different strands completes the library for sequencing.^8,9,12^

Here, we demonstrate the performance of this in-solution OS-Seq approach using a variety of reference DNA samples. We further show its broader applicability to clinical samples, including FFPE biopsies. Using a 130-gene panel, we confirm the technical reproducibility and high performance of the in-solution OS-Seq assay in terms of on-target coverage, uniformity and ability to detect single nucleotide variants (**SNV**s), insertions and deletions (**indels**) and copy number alterations (**CNAs**) from as little as 10 ng of input DNA.

## RESULTS

### Overview of in-solution OS-Seq

The in-solution version of OS-Seq involves three general steps **(Figure 1)**. First, the assay uses a repair process wherein damaged bases present in genomic DNA isolated from FFPE samples are removed by excision only, without implementing a corrective repair step. Next, the DNA sample is fully denatured to single-stranded DNA followed by single-stranded ligation of the adapter. This approach ensures that all DNA species, whether present in single-stranded or double-stranded form, can be interrogated, regardless of starting material quality and quantity. Moreover, optimization of this ligation reaction to a high conversion rate eliminates the need for a whole genome amplification step. Finally, the enrichment of the desired genomic targets – “capture” – occurs with massively multiplexed pools of target-specific primer-probe oligonucleotides that are designed to tile across both strands of the regions of interest at high density; on average, one primer per every 70 bp. Following hybridization, the primer provides a start site for polymerase extension, which captures the targeted DNA molecule, thus completing the library for sequencing through incorporation of the second sequencing adapter. The high efficiency of both the ligation and capture steps minimizes the use of PCR following target capture to 15 cycles, irrespective of the input quantity, to expand the library to sufficient quantities for loading onto the sequencer. For paired-end sequencing, the first read (Read 1) is generated from the synthetic target-specific primer-probes and therefore is at a fixed position within the genome. The second read (Read 2) is generated from the universal adaptor and initiates at a position within the genome corresponding to the 5’-end of the input DNA fragment.

**Figure 1.**
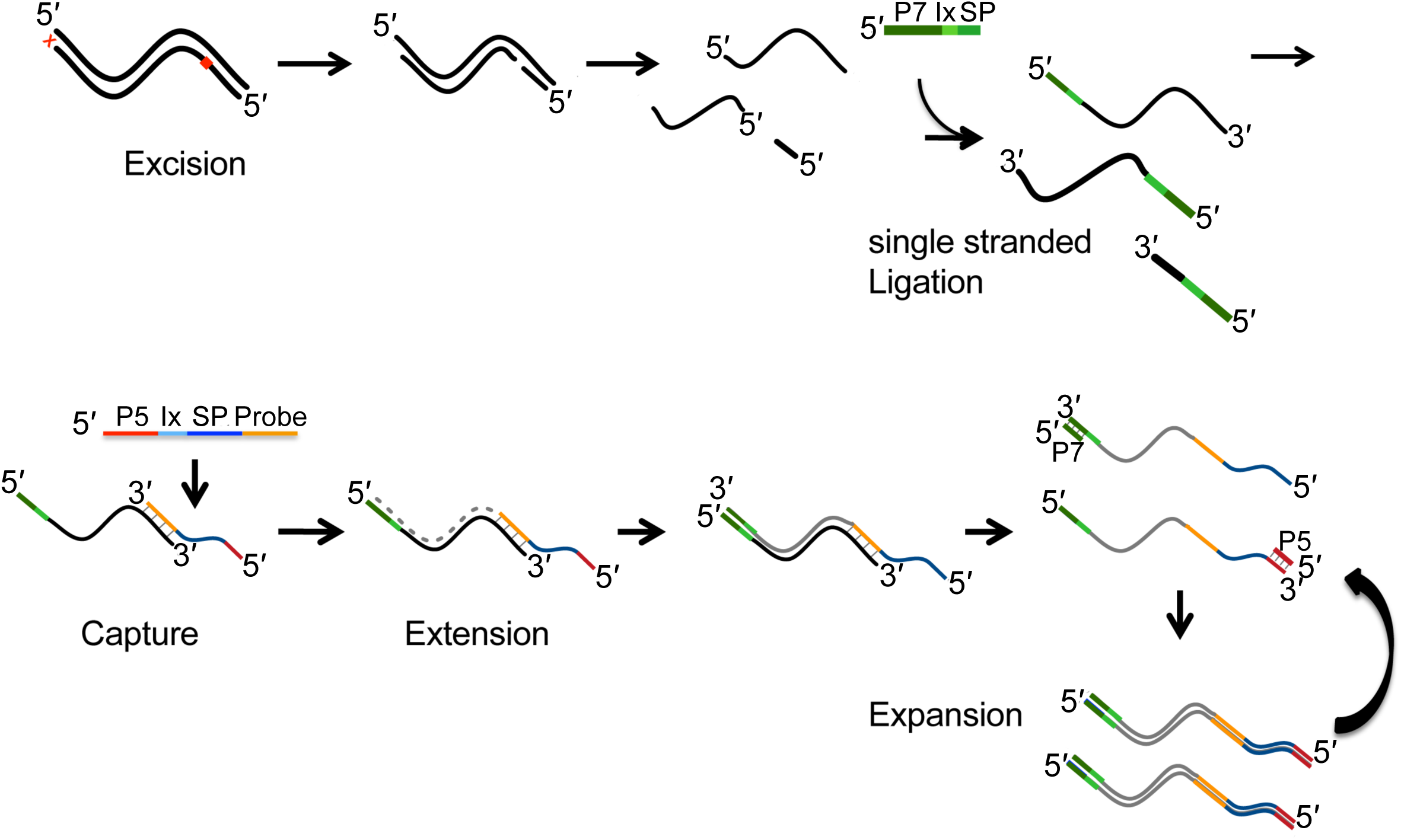
Overview of in-solution OS-Seq process. Damaged bases are removed by excision only, without implementing a corrective repair step The DNA is then denatured followed by adapter ligation to single stranded DNA. In-solution capture using primer-probes is performed for ∼2h, followed immediately by extension to complete the library. Finally, the sequence library is expanded by PCR using primers targeting the P7 and P5 regions to generate sufficient quantities of library for sequencing. 5’ and 3’ ends indicated, P7 and P5 indicate regions of adapters and probes, respectively, required for clustering on the Illumina flow cell, or in the “expansion” section, they indicate PCR primers complementary to the P7 and P5 parts of the adapters and probes, respectively. “Ix” stands for index sequence, and “SP” for sequencing primer-binding site.

We developed a 130-gene panel as an assay for clinical cancer samples **(Supplementary Table 1).** The panel is composed of cancer-related genes, including established tumors suppressors and oncogenes, some of which are known to contain clinically actionable cancer mutations or to provide prognostic information across different malignancies **(Methods)**. All exons of these genes are targeted by primers sets as described in the Methods, making up a set of regions of interest **(ROI)** totaling 419.5 kb.

**Table 1.** Sequencing metrics for control DNA samples

| **Cell line** | **Input DNA (ng)** | **Number of replicates** | **Target coverage, mean (SD)** | **% on target bases, mean (SD)** | **Fold 80 base penalty, mean (SD)** |
| --- | --- | --- | --- | --- | --- |
| NA12878 | 300 | 4 | 3,097 (125) | 85% (0%) | 1.77 (0.01) |
|  | 100 | 4 | 3,028 (149) | 79% (0%) | 1.96 (0.01) |
|  | 30 | 4 | 2,342 (161) | 78% (1%) | 2.20 (0.04) |
|  | 10 | 4 | 2,735 (289) | 67% (3%) | 3.57 (0.33) |
| HD753 | 100 | 3 | 6,941 (739) | 56% (1%) | 1.85 (0.01) |
|  | 30 | 3 | 3,920 (301) | 56% (1%) | 1.87 (0.08) |
|  | 10 | 3 | 4,045 (727) | 51% (2%) | 2.22 (0.16) |
| HD200 | 300 | 4 | 4,441 (1,312) | 73% (1%) | 1.96 (0.05) |
|  | 100 | 4 | 4,509 (1,073) | 66% (1%) | 2.05 (0.10) |
|  | 30 | 4 | 2,766 (507) | 62% (1%) | 2.29 (0.06) |
|  | 10 | 4 | 1,721 (214) | 61% (1%) | 2.64 (0.11) |

SD: standard deviation

### Analysis of the reference genome NA12878

As a preliminary assessment of assay performance, we conducted a mass titration experiment using the Coriell Institute DNA sample NA12878. The genome of this sample has been sequenced extensively under a variety of NGS platforms: it has been used as a pilot genome for the Genome-in-a-Bottle **(GIAB)** consortium, and a genomic reference material from the National Institute of Standards (**NIST**). Because the availability of a high confidence list of ground truth variants for this genome developed by GIAB, NA12878 is widely used for assessment of germline variant detection accuracy.^13^ We performed four independent technical replicates of the assay across DNA inputs of 300 ng, 100 ng, 30 ng and 10 ng. Following library quantification with ddPCR, the same number of molecules across these libraries were pooled and sequenced.

Analysis of the sequencing results showed high on-target coverage across all samples regardless of DNA input quantity. At 300 ng of input DNA, we observed a mean on-target average coverage of 3,097X ± 125 across all four technical replicates. The depth of coverage was maintained even at 10 ng of DNA input, where the mean on-target coverage was 2,700X ± 289 **(Table 1)**. The fraction of on-target reads (i.e. the fraction of reads originating from properly placed primers) was also high regardless of the starting amount of DNA. At 300 ng input DNA, we observed an on-target read fraction of 85% across all replicates with no discernible variance. More significantly, at an input quantity of 10 ng, the on-target read fraction was still high at 67% ± 3 across all replicates **(Table 1, Supplementary Table 2, Supplementary Figure 1a)**.

**Table 2.**
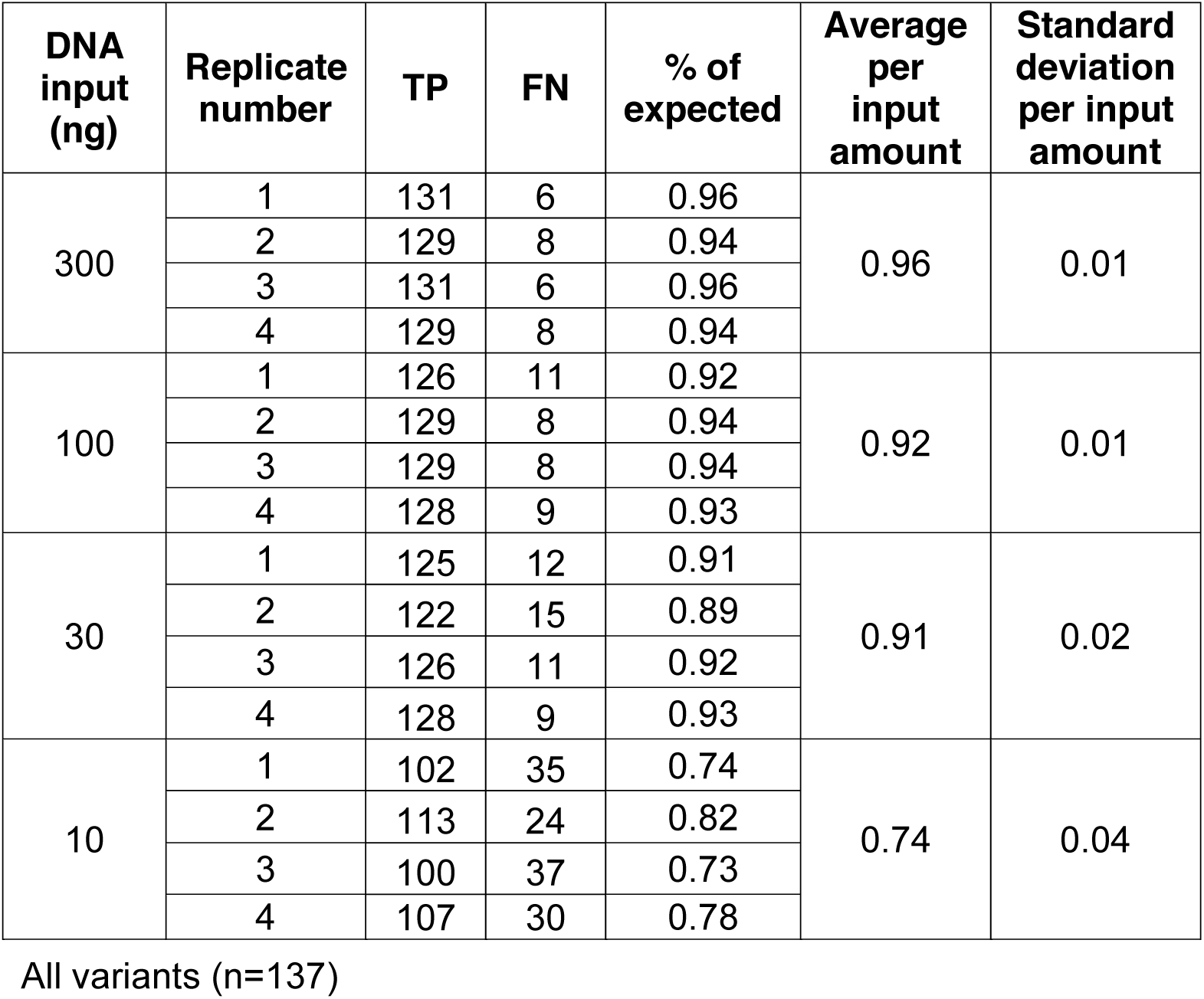
Detection of SNV and indel variants from NA12878

Read coverage uniformity across ROIs was assessed using the fold 80 base penalty metric^14^ and by observing the fraction of ROI bases achieving a set of coverage thresholds ranging from 2X to 100X. The fold 80 base penalty is defined as the fold change of read coverage needed to bring 80% of the ROI bases to the observed mean coverage. A lower value indicates less variability among the coverage of the individual targets; a hypothetical case of perfect uniformity would have a fold 80 base penalty of 1. We noted that a high level of uniformity was achieved across the range of input quantities, with fold 80 base penalty values ranging from 1.77 (SD=0.01) for 300 ng of input DNA to 3.57 (SD=0.33) for 10 ng **(Table 1 and Supplementary Table 2, Supplementary Figure 1b)**. This compares favorably with published high quality exome sequence data sets, where the fold 80 base penalty typically ranges between 2 to 4 ^15–17^. In these exome datasets, the lower fold 80 base penalties are generally achieved using microgram-range amounts of input DNA^15–17^, which is in contrast to our results using nanogram-range input quantities. Importantly, the uniformity in coverage resulted in a high fraction of targeted ROI bases being covered at read coverage of 100X or more, with 98% covered at 300 ng, and 92% at 10 ng **(Supplementary Table 2 and Supplementary Figure 1c).**

To determine the assay’s performance in detecting germline variants, we investigated its ability to detect the ground truth variants present in the intersection of the GIAB high-confidence regions with the ROIs interrogated in our assay. This intersection includes at total of 137 variants distributed among 128 SNVs, and 9 indels. At the highest input quantity of DNA (300 ng), we determined that 96 ± 1% of GIAB ground truth variants were detected in our unfiltered calls. With 10-fold less material (30 ng), 91 ± 2% of GIAB reference variants were detected (**Table 2, Supplementary Figure 2,**).

**Figure 2.**
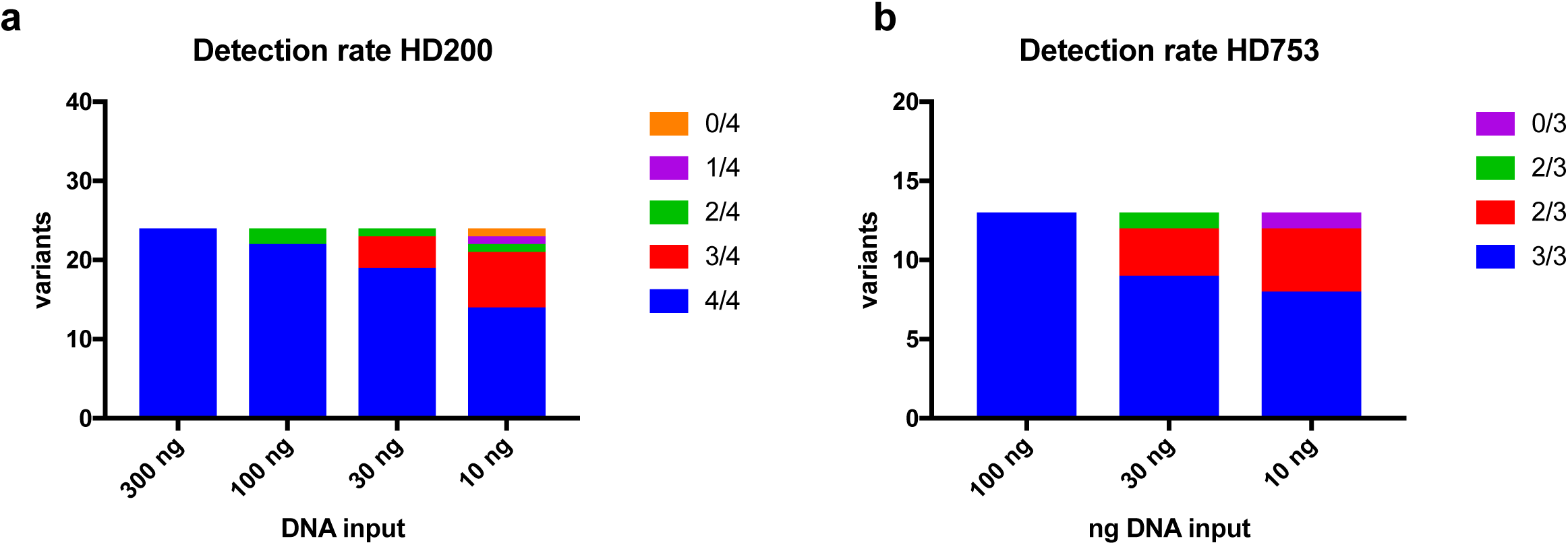
Analysis of variant allelic fraction. a) Detection rate of spiked-in somatic variants in HD200. Variants detected in 4 out of 4 replicates are shown in blue, 3 out of 4 in red, 2 out of 4 in green, 1 out of 4 in purple, and 0 out of 4 in yellow. b) Detection rate of spiked-in somatic variants in HD753. Variants detected in 3 out of 3 replicates are shown in blue, 2 out of 3 in red, 1 out of 3 in green, and 0 out of 3 in purple.

### Detection of variants at different variant allelic fractions

We assessed the ability of the 130-gene assay to detect variants present at different VAFs, mimicking the distribution of VAFs expected for somatic mutations in tumor tissue samples. We used a set of reference materials derived from either mixtures of engineered cell lines or synthetic DNAs harboring specific variants spiked into a reference background genome. Either approach yields DNA samples with well-known somatic mutations at pre-validated allelic fractions. First, we analyzed the STMM-Mix-II reference standard, which includes 37 known cancer somatic variants (24 SNVs and 13 indels) within the ROIs covered by our assay (**Supplementary Table 3**). These variants are spiked-in at known VAFs within the background of the NA24385 genome, another GIAB analyzed genome and NIST reference material. We obtained a dilution series of STMM-Mix-II at 5, 10, 15, and 25% VAFs for the 37 mutations, with the VAF for each somatic variant validated with droplet digital PCR **(ddPCR)** by the manufacturer. Based on the GIAB list of high confidence ground truth germline variants for the genome of NA24385, we determined whether detected variants are either somatic, germline, or false positive events. This allowed us to calculate sensitivity and specificity for SNV/indel detection.

**Table 3.**
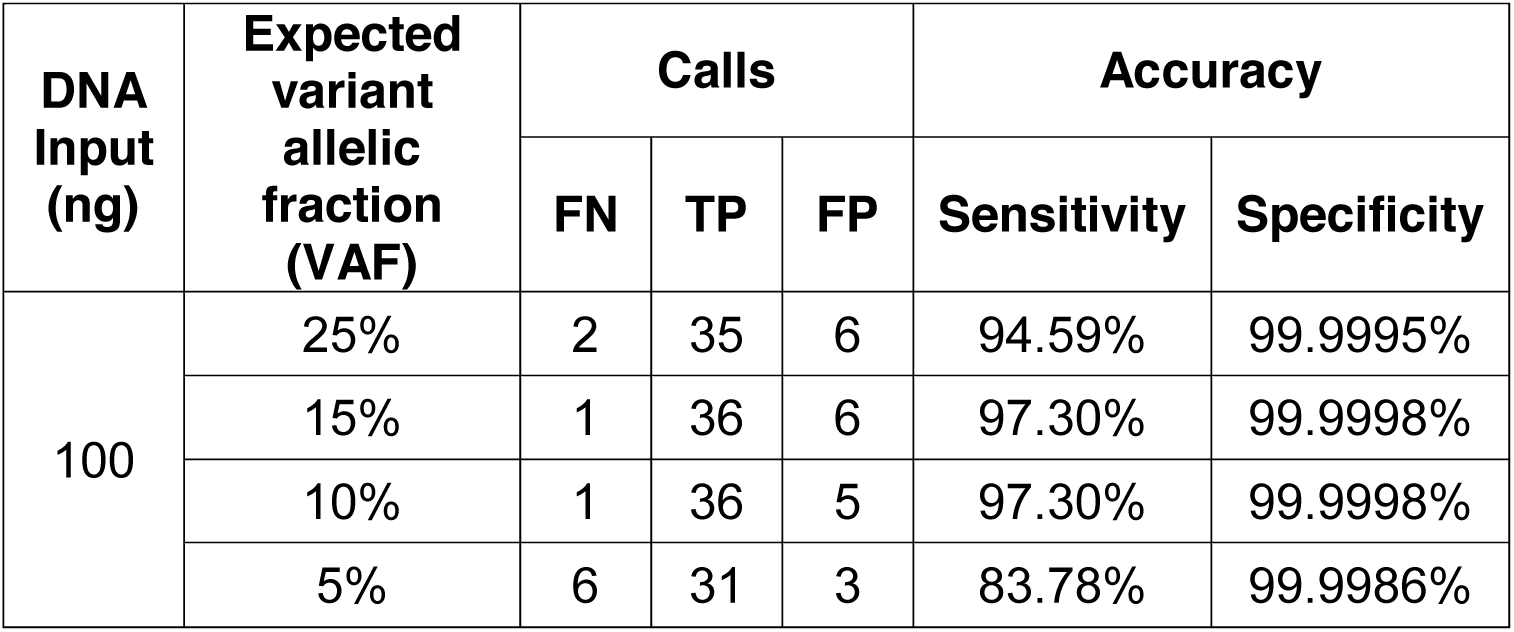
Detection of variants from control DNA mixtures

| DNA Input (ng) | Expected variant allelic fraction (VAF) | Calls |  |  | Accuracy |  |
| --- | --- | --- | --- | --- | --- | --- |
|  |  | FN | TP | FP | Sensitivity | Specificity |
| 100 | 25% | 2 | 35 | 6 | 94.59% | 99.9995% |
|  | 15% | 1 | 36 | 6 | 97.30% | 99.9998% |
|  | 10% | 1 | 36 | 5 | 97.30% | 99.9998% |
|  | 5% | 6 | 31 | 3 | 83.78% | 99.9986% |

From each DNA mixture, we used 100 ng input DNA. Overall, the sequencing metrics for each DNA mixture were comparable to that observed with NA12878, with on-target average coverage being greater than 1,690X across all samples and replicates **(Supplementary Table 2)**. The assay demonstrated a specificity of ∼100% regardless of the sample’s VAF (**Table 3**). Sensitivity was consistently high with VAFs at 10% or greater having a sensitivity of more than 90.0%. At a VAF of 5%, a sensitivity of 83.8% was observed, indicating the general high performance of the assay for low allelic fractions.

To test the performance of the assay on DNA of compromised quality, we relied upon the HD200 FFPE reference material. This reference material consists of a mixture of the colorectal cancer cell lines HCT116, RKO and SW48 at defined ratios and includes frequently occurring cancer mutations at VAFs lower than 50% validated with ddPCR by the manufacturer. Further, the sample has been subject to FFPE processing as a surrogate for archival tissue. Twenty-four of the nonsynonymous mutations within this sample are covered in the 130-gene assay ROI. Overall, we obtained sequencing metrics similar to what was observed with NA12878 **(Table 1** and **Supplementary Table 2)**. Average on-target coverage ranged from 4,509X ± 1,312 at 100 ng to 1,721X ± 214 for 10 ng input DNA. The fraction of on-target reads was greater than 50% regardless of the amount of input DNA across replicates. Moreover, the average fold 80 base penalty ranged from 1.85 at 100 ng to 2.22 at 10 ng. This was slightly better than observed for NA12878, underlining the consistency and uniformity of coverage at the lowest input amount of FFPE DNA. In aggregate, 82% of variants were detected in four out of four replicates, and 94% were found in at least three replicates **(Figure 2 a**, **Supplementary Table 4)**. As the genetic background of the cell lines used in the construction of this reference material is not well characterized, we were unable to calculate specificity from this data.

**Table 4.** CNA calling from a control DNA sample

| Amount of DNA (ng) | Gene | Read depth CNA calling |  | Varscan2 CNA calling |  |
| --- | --- | --- | --- | --- | --- |
|  |  | Expected ratio* | Observed ratio, mean (SD) | Expected ratio (log scale)* | Observed ratio (log scale), mean (SD) |
| 100 | *MYC* | 4.90 | 4.09 (0.04) | 2.29 | 2.51 (0.15) |
|  | *MET* | 2.25 | 1.87 (0.04) | 1.17 | 1.24 (0.14) |
|  | *ALK* | 1.32 | 1.41 (0.04) | 0.40 | 0.66 (0.14) |
| 30 | *MYC* | 4.90 | 3.67 (0.05) | 2.29 | 2.93 (0.08) |
|  | *MET* | 2.25 | 1.51 (0.03) | 1.17 | 1.61 (0.08) |
|  | *ALK* | 1.32 | 1.40 (0.03) | 0.40 | 1.11 (0.09) |
| 10 | *MYC* | 4.90 | 4.52 (0.24) | 2.29 | 3.26 (0.11) |
|  | *MET* | 2.25 | 1.64 (0.07) | 1.17 | 1.49 (0.20) |
|  | *ALK* | 1.32 | 1.49 (0.04) | 0.40 | 1.16 (0.20) |

SD: standard deviation

Next, we used the reference material HD753 to test the performance of the assay in identifying copy number alterations in addition to somatic SNVs/indels. The HD753 DNA contains validated copy number alterations, engineered into the genomes of a set of background cell lines in addition to 18 validated cancer somatic mutations. Of these 18 somatic variants, 13 overlap with the ROIs within the 130-gene assay **(Supplementary Table 5, Figure 2 b)**. Sequencing metrics obtained from input quantities ranging from 100 to 10 ng were found to be equivalent to those observed in other samples analyzed **(Table 1** and **Supplementary Table 2)**. Across the entire range of input DNA, 78% of SNVs and indels were found in all three replicates, and 95% were found in at least two replicates **(Supplementary Table 5, Figure 2 b)**. Even at 10 ng of input DNA, we detected all variants with the sole exception of an insertion mutation in *EGFR* (V769D770insASV). As in the case of HD200, the lack of detailed characterization of the genetic background of the cell lines underlying HD753 did not allow the calculation of the specificity of SNV/indel detection for this reference sample.

Finally, as the HD753 sample harbors two previously characterized amplifications in the *MET* and *MYC* cancer drivers, both of which are present in the 130-gene assay, we assessed the performance of the 130-gene panel in identifying CNAs. A range of DNA input amounts including 100 ng, 30 ng, and 10 ng were tested across three technical replicates using NA12878 as a normal diploid DNA control (**Figure 3, Table 4**). We used Varscan2^18^ and a custom method that identified outliers in the log_2_ ratios of the median coverage depth across all ROIs between the test and negative control samples to determine CNA values **(Methods)**. Both *MYC* and *MET* amplifications were identified by both methods at the expected ratios and across all the input amounts tested. Additionally, an *ALK* gene amplification was identified that was not previously reported in this material **(Figure 3)**. Commercial ddPCR CNA assays verified this amplification, confirming both the presence and magnitude of the *ALK* amplification as determined by OS-Seq **(Table 4)**.

**Figure 3.**
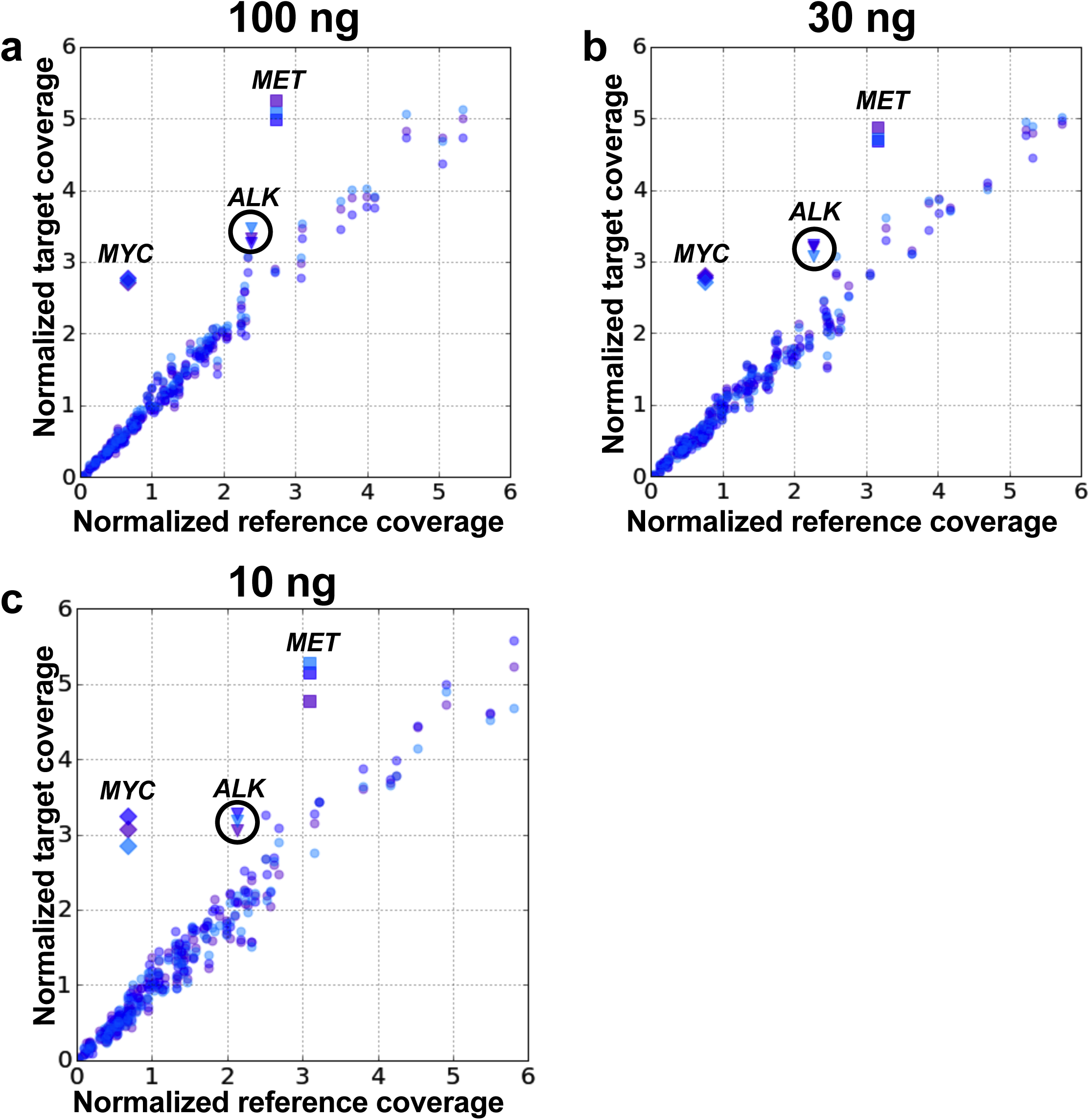
Detection of copy number alterations. Normalized coverage for all genes in the 130-gene panel for each replicate of HD753 (target) plotted vs normalized coverage for all genes in NA12878 (control, the same control is used for comparison with each target replicate) for 100 ng (a), 30 ng (b) and 10 ng (c) DNA input. Each replicate is shown in a different color. The three amplified genes are shown as diamonds (*MYC*), squares (*MET*) and triangles (*ALK*).

### Analysis of clinical FFPE tumor and matched normal DNA

Given the observed performance on the above reference materials, we evaluated the assay’s ability to detect variants from DNA extracted from a variety of clinical samples. Commercially sourced patient-matched blood and tissue samples from Stage III lung and colorectal patients were used **(Supplementary Table 6)**. Cell-free DNA (**cfDNA**) was isolated from plasma, as well as genomic DNA both peripheral monocytes and archival FFPE tumor tissue. Following repair, 100 ng of FFPE and PBMC derived DNA, and 40 ng of cfDNA, was input to adapter ligation. We used DNA extracted from PBMCs (a high quality DNA source) to compare sequencing quality metrics with those obtained from the FFPE-extracted DNA.

Based on the metrics described above, our assay exhibited similarly robust performance on both PBMC and FFPE samples compared to the performance observed with high quality genomic DNA extracted from cell lines **(Supplementary Table 2)**. On-target coverage ranged from 2,300X to 5,600X. The fraction of on-target reads was consistently high at greater than 50%, regardless of input DNA (PBMCs or FFPE).

We examined the assay’s ability to identify variants in these clinical samples, beginning with germline variants from the matched pairs of PBMC and FFPE **(Figure 4)**. To assess the quality of the germline genotypes identified, we compared our data with the database of common germline variants developed by the Exome Aggregation Consortium **(ExAC)** and the 1000 Genome Project.^19,20^ Greater than 90% of the SNV calls from the PBMC and FFPE samples were also reported in the ExAC and 1000 Genome Projects **(Supplementary Table 7)**, indicating that the quality of these calls is very high: most true positive germline variants are expected to have been previously identified in these projects, which have extensively catalogued common genetic variation across major populations. We also compared the SNV overlap between the calls obtained from matched PBMC and FFPE tissues DNA. As we noted, FFPE DNA is chemically modified in ways that can impair sequencing data quality, which could lead to compromised variant calling. We observed a large overlap of germline variants called in patient-matched FFPE and PBMCs samples, ranging from 79 to 91% of variants being found in both tissue samples. This result indicates that the quality of our calls is high and not overly compromised by the tissue source.

**Figure 4.**
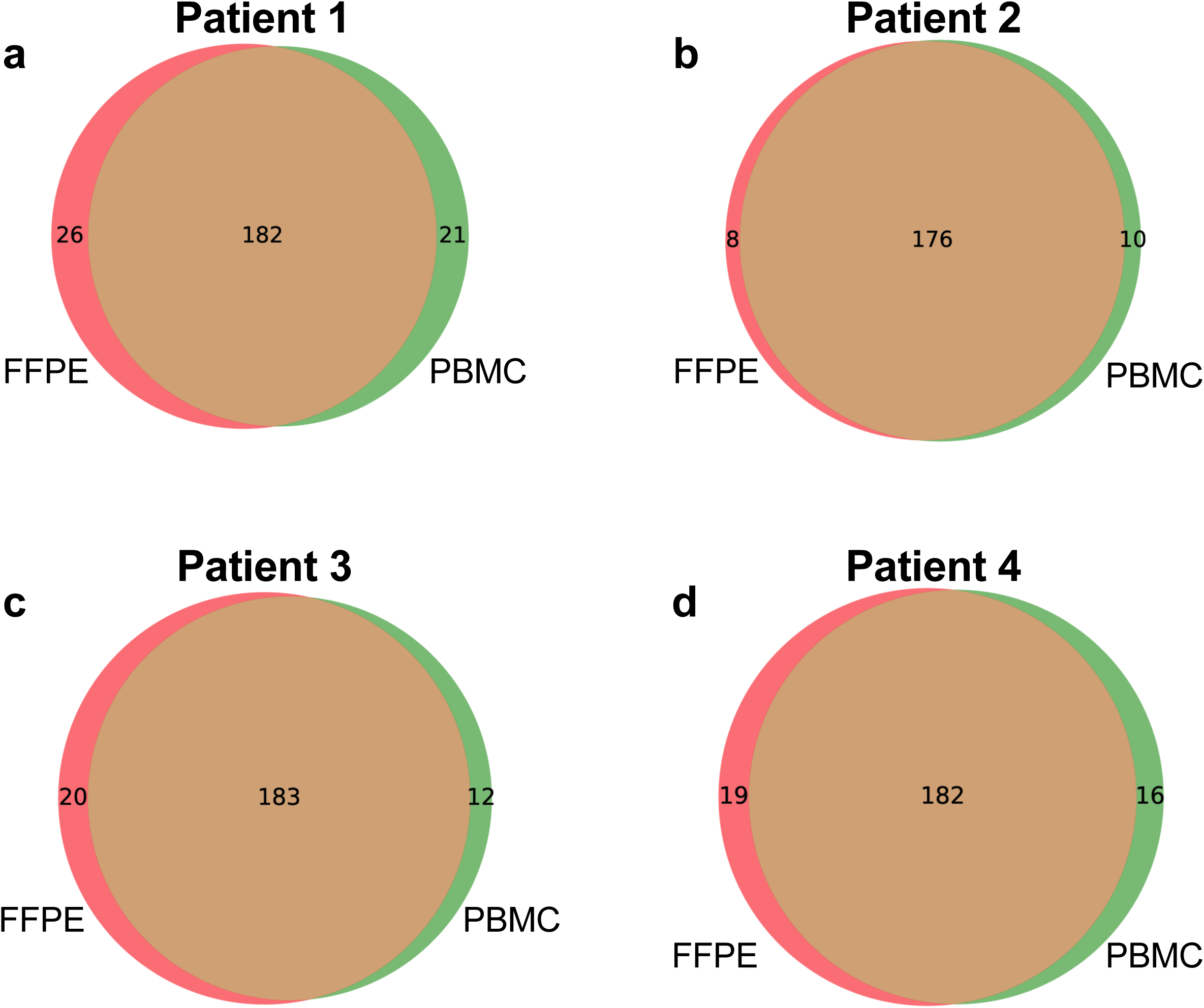
Variant overlap between DNA from PBMCs versus FFPE tissue. Overlap of variants called by GATK in the FFPE (red circles) and PBMC (green circles) samples from the four patients included in the matched sample study. a-d shows Patient 1-4, respectively.

Somatic mutations within the FFPE sample were identified using the patient-matched normal DNA derived from PBMCs.^21^ The somatic mutations observed are reported in **Supplementary Table 8** along with a variety of annotations from the COSMIC database, including the reported frequency of occurrence of these mutations in colorectal and lung cancers.^22^ From all four samples, we identified a total of 32 mutations from the 130 genes that had a high likelihood of being pathogenic, had multiple read support and occurred above a general overall depth threshold of 40X. Of the detected somatic variants, 15.6% had previously been reported in other tumors. In Patient 1’s colorectal cancer, we detected a stop codon in *APC,* a well-documented cancer driver mutation in an essential tumor suppressor gene involved in colorectal cancer oncogenesis.^23^ We also identified mutations in *ERBB2* (i.e. HER2), which have recently been found to be mutated in CRC and may represent a gene for targeted therapy in this cancer.^24^ Finally, we found mutations in *NF1*, also reported mutated previously in CRC.^25^

Patients 2, 3 and 4 were diagnosed with non-small cell lung cancer **(NSCLC)**. For Patient 2, we discovered a mutation in *CDKN2A*, frequently inactivated in lung cancer.^26^ Patients 3 and 4 both exhibited mutations - distinct for each patient - in *FGFR3,* reported mutated with a frequency of up to 3% in NSCLC.^26^ Patient 4 also had a mutation in *ABL1*, reported mutated with a frequency of 1.5% of NSCLC.^27^ None of the patients had overlapping mutations.

As a final proof of concept, matched cfDNA samples from each patient were sequenced using the OS-Seq protocol. Sequencing metrics were equivalent to those found in both reference DNA and genomic DNA from clinical samples (**Supplementary Table 2**). The assay’s ability to detect overlapping germline variants in cfDNA was determined in both PBMCs and FFPE samples **(Supplementary Figure 3)**. We found a high proportion of cfDNA variants also observed in PBMC and FFPE samples: 75-91% of cfDNA variants were called in all three sample types, with 79 to 94% were found in at least one additional sample type.

In addition to performing the matched sample comparisons described above, we also investigated the assay’s ability to remove common FFPE-derived false positive variants, which most frequently arise from cytosine deamination to uracil, resulting in an observed C to T transition (G to A on the opposite strand) ^28^. Dexyuridine, which is the result of cytosine deamination in DNA, can be removed enzymatically ^29^. Such removal is typically included in FFPE repair processes, including the excision step used in our protocol. To investigate whether this process works efficiently, we subjected four FFPE samples (One of the matched FFPE samples already sequenced and three additional commercially available samples) to either our standard excision process or no repair. We also included another enzymatic repair process for FFPE DNA as an additional control **(Methods)**. We processed these samples through our standard library preparation method starting after repair, and sequenced the resulting libraries. We analyzed the variant data. When we compared the ratio of C to T and A to G (C>T/A>G) variants between the different repair processes, we found that our standard excision and the alternative FFPE repair both generated C>T/A>G ratios of 37-58%, and were very similar to each other (differing by 0-3 percentage points) for each individual sample. In the samples not treated with repair, the C>T/A>G ratios were 13-20 percentage points higher than the treated samples, with a ratio of 51-71% (**Supplementary Table S9**). These results demonstrate that the assay’s excision step efficiently removes FFPE induced damage and compares well to other commercially available repair process kits. Importantly, the comparison to the no repair samples demonstrated that the excision step significantly contributes to the removal of nucleotide modifications resulting from FFPE chemical processing.

## DISCUSSION

The sequencing of tumor tissue samples derived from clinical biopsies presents several major challenges. First, tumor tissues are a complex mixture of adjacent normal cells and potentially multiple clones of cancer cells. As tumor purity in clinical specimens can be as low as 20%, deep sequencing is required to detect somatic mutations present in lower allelic fractions. This factos has motivated the use of targeted sequencing of actionable cancer genes for improved sensitivity. Second, most readily available clinical tumor samples are archived in FFPE blocks, a process aimed at enabling histological evaluation that results in damage to DNA. Third, abundance of nucleic acid analyte in clinical samples is frequently low, compromising the performance of NGS-based assays. Finally, genetic biomarkers that can inform therapy decision and/or prognosis include CNAs in addition to SNV and indels. Unfortunately, CNAs constitute a type of somatic mutation that is more difficult to detect with commonly used targeted assays.^30^.

The clinical implementation of targeted sequencing assays commonly involves positive selection of ROIs, either through PCR amplification (amplicon-based approaches) or through hybridization enrichment with long oligonucleotides (bait hybridization).^12^ However, both approaches are ill-suited to addressing the challenges associated with processing FFPE clinical samples. Amplicon-based targeting approaches require that both forward and reverse primers hybridize to the target, and thereby capture the ROI. This is particularly challenging with fragmented DNA from FFPE clinical samples. Optimization requires either a reduction in the amplicon footprint to increase the likelihood that both primers are able to hybridize, and/or requiring extensive amplification to generate sufficient amounts of sequencing library material from the fraction of molecules upon which both primers can hybridize. Moreover, in the presence of damaged bases, the amplification efficiency of PCR is reduced when using DNA polymerases with proof-reading capability, as these types of enzymes have poor tolerance for base modifications such as those generated through deoxy-cytosine deamination to deoxyuracil or through depurination to abasic sites typically present in FFPE.^6^

In contrast, bait hybridization based approaches mitigate the need for two primers to capture a ROI by instead capturing any fragment within the ROI that can hybridize to the bait oligonucleotide. However, more extensive enzymatic and technical processing of the material is required resulting in a complex workflow and intricacies of preparation that are more prone to experimental error. In particular, following traditional library preparation methods, blunt DNA ends are required in preparation for the initial ligation of double stranded adaptors to the fragmented DNA. This is more problematic for FFPE tissue samples, as the extracted DNA can not only have a high proportion of damaged bases, but also a high proportion of single-stranded molecules that are effectively eliminated from traditional sequencing library preparations^6^. Furthermore, the use of long oligonucleotides to capture ROIs through hybridization increases the likelihood of capturing off-target regions^31^, and requires extensive washing to ensure specificity.^32^ This requirement for stringent washing increases the differential efficiency in retention of regions of varying GC-content, affecting the uniformity of coverage across genomic targets and potentially resulting in false negative results.^33^

Both the amplicon-based and the bait hybridization approaches require extensive PCR amplification to generate a sufficient number of sequencing library molecules, as well as to mitigate negative factors due to sample quality and/or elaborate processing steps. Extensive amplification exacerbates biases associated with GC-content and length, which can skew the representation of the original DNA templates within the sequenced library. ^34^ This skewing results in reduced sensitivity to the identification of genetic alterations such as copy number variants.^35,36^ Overall, these issues affect the diagnostic yield of NGS targeted assays.

To address these issues, we developed an in-solution targeted sequencing assay based on an enzymology significantly different to previously described methods. This method relies on primer annealing and extension of single stranded DNA.^8,9^ As we have demonstrated here, our in-solution OS-Seq assay has been optimized for high performance on low input quantities and compromised nucleic acid quality from clinical specimens. Specifically, in-solution OS-Seq enables sampling of degraded and fragmented single stranded DNA with high efficiency and on-target rates while utilizing limited PCR amplification. Consequently, the assay demonstrates high uniformity coverage at DNA inputs down as low as 10 ng of DNA, enabling an efficient means of performing deep sequencing of target regions with low incidence of false negatives for somatic and germline variants.

As another added feature, the combination of optimized hybridization conditions and dense tiling across both DNA strands by extension primers delivers high performance, as measured by a low fold 80 base penalty metric values and a high fraction of ROI bases covered at ≥ 100X, compared to other assays.^15,17,21^ As a result, there are fewer genomic regions of interest with low read coverage where false negatives can occur, increasing resilience when analyzing poor samples with limited DNA inputs. Further, such uniform coverage enables cost-efficient lab operations without the need to compensate for low coverage regions by increasing coverage through excessive sequencing.

A high performance NGS assay should detect the mix of variants often present at low VAF within a tumor sample. When validating clinical tests for tumor sequencing, there has been significant emphasis on controlling the false positive rate with little examination for false negatives.^37–39^ In addition, assay validation is typically done using diploid cell lines that are not cancer – these lines do not provide a broad allelic distribution of somatic variants seen in cancer tissues.^39^ Consequently, accurate estimates of sensitivity and specificity of targeted resequencing assays for tumor profiling become difficult to perform.^40,41^ In our current study, we have addressed this challenge by using a combination of reference materials, including recently available standards with well-characterized genomes in the background of somatic variants spiked in at varying % VAFs.

The OS-Seq 130-gene panel presented here covers a comprehensive set of actionable cancer genes, generating a breadth of information that permits clinicians to identify somatic mutations linked to approved, off-label, and investigational drugs,^42^. We demonstrate high performance on low quality input material and provides both high and uniform coverage, which allows the identification of clonal and sub-clonal somatic mutations even from low cellularity tumors. For these reasons, the OS-Seq assay is particularly well suited for the poor quality of real clinical specimens, and will increase the yield of clinically actionable variants to inform prognosis and cancer therapy selection.

## MATERIALS AND METHODS

### DNA samples and preparation

Purified genomic DNA **(gDNA)** from the NA12878 Coriell cell line was obtained from the Coriell Institute for Medical Research (Camden, NJ). Purified DNA from the structural multiplex reference standard HD753 was obtained from Horizon Diagnostics (Cambridge, UK). The SeraCare STMM-Mix-II standard was acquired from SeraCare (Milford, MA). Curls of FFPE cell line mixtures (HD200) with defined allelic frequencies were obtained from Horizon Diagnostics (Cambridge, UK). Anonymous matched plasma, buffy coat and FFPE solid tumor samples from stage III lung and colorectal cancer patients were purchased from Indivumed GmbH (Hamburg, Germany). Blood components were shipped on dry ice and stored at -80 °C until ready for processing.

The genomic DNA was purified from two 10-20 μm FFPE curls using the ReliaPrep FFPE gDNA Miniprep System (Promega, Sunnyvale, CA), with the following modifications: FFPE curls were incubated for 16 hrs overnight with proteinase K at 65 ° C in lysis buffer. Following a 1h incubation at 90 ° C, tubes were flash cooled, and the entire mixture transferred to a microfiltration device equipped with a 0.45 μm cellulose acetate filter (Corning COSTAR, Corning, NY). Upon centrifugation for 15 min at 4 ° C at 16,000 × *g* to remove particulates, the filtrate was processed according the manufacturer’s guidelines.

Buffy coat samples were gently resuspended in 500 μL phosphate-buffered saline and transferred to a 15 mL conical tube. Residual red blood cells were then lysed by the addition of 4.5 mL of ACK lysis buffer (ThermoFisher Scientific, Carlsbad, CA) and incubation for 10 minutes with inversion at room temperature. Peripheral blood mononuclear cells (PBMCs) were then pelleted via centrifugation for 10 min at 1,600 × *g*. Pelleted cells were then resuspended in 400 μL of cell lysis buffer (50 mM Tris·HCl, 50 mM Na·EDTA, 0.1% Triton-X100 1.0% sodium dodecyl sulfate, pH 8.0) with 20 μL of >600 mAU/mL proteinase K (Qiagen) and 20 μL of 100 mg/ml RNase A (Qiagen). Following incubation for 1 hr at 65°C, ∼0.7 volumes (350 μL) of neat isopropanol was added and the solution mixed by gentle inversion. After incubation for 30 min at -20°C, samples were centrifuged at 16,000× *g* for 15 min, and the supernatant removed. Pellets containing genomic DNA were then washed once with 1 mL of freshly prepared 70% Ethanol, and air-dried for 5 min at room temperature, followed by resuspension in 300 μL IDTE buffer (Integrated DNA Technologies, Coralville, IA).

The cfDNA was purified from 3 mLs of plasma using the QIAamp Circulating Nucleic Acid Kit (Qiagen, Redwood City, CA) according to the manufacturer’s recommended guidelines. All samples, with the exception of cfDNA samples, were mechanically sheared prior to input into to the TOMA OS-Seq protocol. Briefly, up to 1 μg of DNA was sheared either with a Covaris E210R (Covaris, Woburn, MA) or a ST30 (Microsonic Systems, San Jose, CA) sonicator to a target base pair peak of 600 bp according to the manufacturers’ recommendations.

### DNA quantification

DNA samples were quantified with ddPCR using the *RPP30* gene as a surrogate for the number of genomic equivalents. For each sample to be analyzed, ddPCR reactions were prepared using 11 μl of Droplet PCR Supermix for probes, 1.1 μl of HEX-labeled PrimePCR^™^ ddPCR^™^ Copy Number Assay: RPP30, Human (Assay ID: dHsaCP2500313; BioRad, Hercules, CA), 2.2 μl gDNA, and nuclease free water to a final volume of 22 μl. 20 uL of this reaction mixture was then processed and analyzed on the QX200^™^ Droplet Digital^™^ PCR System according to the manufacturer’s recommended guidelines using QuantaSoft v1.7.4.0917 (BioRad, Hercules, CA). Values were converted from copies/μL to ng/μL using 30 ng per 10,000 copies of genome equivalents.

### Targeting assay

The TOMA COMPASS 130-gene kit (TOMA Biosciences, Foster City, CA) includes a set of 14,050 OS-Seq primers designed to cover 2,111 ROIs encompassing the exons of 130 genes. Briefly, to select the set of targeting sequences, a melting temperature compatible with the annealing temperature was selected to delineate candidate primers considering the annealing buffer composition, and sequences were scored with an empirical scheme that accounted for both intrinsic features of the primer sequence, such as G+C content, homopolymers, and secondary structure, as well as genomic features such as the presence of SNPs identified within the dbSNP database, relative target position, the anticipated contribution to ROI coverage, and the predicted specificity of the primer across the genome. Finally, potential interactions between primers in the same pool were evaluated. After evaluation, candidate sequences with scores below a threshold were discarded, and the highest scoring sequences were selected to target each ROI.

Samples were processed using the TOMA COMPASS 130-gene library preparation kit according to manufacturer’s recommendation (TOMA Biosciences, Foster City, CA). First, up to 1 μg of DNA was used for the TOMA repair. In the cases where no repair was used, the repair buffer was replaced with the same volume of elution buffer, and samples were processed according to the protocol. In the cases where the TOMA repair was replaced by NEB repair (NEBNext FFPE DNA Repair Mix, NEB, Ipswich, MA), the same amount of DNA as for other repair treatments were repaired using the NEB kit following the manufacturer’s recommendations. The repaired DNA was then purified using the TOMA purification protocol using 2 volumes of sREP + beads and resuspended in 40 μl elution buffer. After DNA repair, DNA concentrations were measured via ddPCR as described and an appropriate amount of DNA was used as input to ligation. Adapter ligation, target capture, and library expansion were then carried out according to the TOMA COMPASS 130-gene library preparation kit. A series of 100-fold dilutions of the resulting libraries were performed in TE buffer and the 10^−6^ dilutions were then quantified via ddPCR using the TOMA ILQ assay, using the following PCR cycling parameters: 95°C 10 min; 30 s at 94°C, 30 s at 55°C, 60 s at 70°C, 40 cycles; followed by 5 min at 70°C. The TOMA ILQ assay measures P7 (labeled by FAM) and P5 (labeled by HEX). Linkage values determined through QuantaSoft v1.7.4.0917 identifies the presence of library molecules with both P5 and P7 adapter arms, were used to calculate the number of library fragments per μl.

Based on the library quantification results, 1.0 to 1.4 billion total library fragments were loaded onto the NextSeq 500 (Illumina, San Diego, CA) according to the manufacturer’s recommendations with the following adjustments. Briefly, libraries to be run were pooled, and volume adjusted to 20 μl with TE buffer. The pooled library was denatured by adding 1 μl of freshly prepared 0.5 M NaOH and incubating for 5 minutes at room temperature. Chilled HT1 buffer (1280 μl) was then added to the library and the entire mixture loaded into the Illumina NextSeq 500/550 High Output v2 kit (300 cycle) sequencing cartridge. The sequencing primers were diluted and used as indicated in the TOMA COMPASS 130 library preparation kit protocol. Libraries were then sequenced as paired-ends (2x150 bp).

### Analysis of sequencing data

#### Read Mapping and performance metrics

Before aligning reads, we pre-processed FASTQ files to remove bases where the quality value was less than 28. We used two algorithms for mapping and aligning reads to the human genome reference assembly (hg19 with decoys). We used BWA (v7.1.5) with default settings, or alternatively, we mapped the reads using RTG map v3.7 (Real Time Genomics Ltd., New Zealand). We relied on Samtools^43^ or Picard (Broad Institute, Cambridge, MA) for additional sequence processing and coverage analysis. We identified the OS-Seq primers that generated the read based on a probe metadata file, and tagged the alignment file with the primer. We evaluated paired end reads, and for those sequences with the correct OS-Seq primer sequence we identified the sequences that were located within the ROI targeted by the primer and in correct orientation (plus/minus strand). Sequence reads were called as off-target when they aligned with an insert size larger than 1.5 Kb between sequence read and primer probe.

#### Variant calling in NA12878 and matched samples

For the targeting assay, we created a bed file of target regions using the coordinates of the targeted exons enlarged by an interval of 50 bases on each flank. This file was provided as an input to the variant callers to limit calls to those regions. To eliminate any influence on variant calling from the synthetic primer probe sequences the primer probe bases were removed from the sequence reads prior to variant analysis. For germline variant calling in the NA12878 cell line and PBMC patient samples, we utilized either GATK (v3.4.6) using published best practices^44^ or RTG snp (v3.7) using default parameters. For calling somatic mutations in paired tumor/normal samples, MuTect (v1.1.4) was used with parameters: -rf BadCigar -downsampling_type NONE.^21^ We identified those mutations with multiple read support that were generally seen in regions with 40X coverage. The Combined Annotation-Dependent Depletion **(CADD)** score was used to evaluate variants.^45^ We reported those mutations which had a CADD score greater than 20 and also noted cases where the mutations had been identified in COSMIC.

#### Somatic mutation calling for HD200, HD753, and SeraCare STMM-Mix-II

To call somatic mutations in absence of matched normal sample such as in the case of the reference materials, we used a modification of the Bayesian network variant caller previously described for family pedigrees^46^, describing a tumor/normal network where the tumor node inherits variants from the germline and incurs de novo somatic mutations. In the absence of normal data the germline variants were to be imputed. Germline and somatic priors from the ExAC^19^ and COSMIC^22^ databases were used to score the variants into putative somatic calls. The final VCF files generated were examined for the expected variants. Afterwards, we compiled the sequencing depth and VAF. In addition, the corresponding BAM files were visually inspected and the depth and VAF was recorded. The average and standard deviation of depth and VAF was calculated for each cell line and DNA input amount, and is presented in **Supplementary Table 4** for HD200 and **Supplementary Table 5** for HD753.

#### Benchmarking of variant calls

To evaluate sensitivity and specificity of variant calling with reference materials we compared the test VCF with a ground truth reference VCF using the vcfeval utility of the RTG Tools package (Real Time Genomics Ltd., New Zealand^46^). In the case of the germline calls for NA12878, the ground truth file was the reference dataset released by the GIAB for the high confidence regions that overlap the regions of the 130-gene panel ROIs (v 3.2.2 ^13^). In the case of somatic reference materials (HD200, HD753, SeraCare STMM-Mix-II), we created a synthetic VCF with the corresponding calls as provided by the COSMIC database VCF (v77),^22^ and used vcfeval with the –squash-ploidy option to only consider allele matches. Received operator curves (ROC) were created with the rocplot utility of RTG Tools.

#### Measuring copy number variation

To identify copy number variations we normalized coverage depth of the aligned data across each ROI in the assay by the median across all of the ROIs for the test sample and a negative control diploid cell line (NA12878). We then calculated log_2_ ratios of the test sample and the negative control at the ROI and then at the gene level, to eliminate region specific biases. To establish if a log_2_ ratio value for a given ROI was significantly different from the rest of the population, we applied the Thompson Tau test for outliers (t = 2.629; 2-tailed inverse t-distribution at a = 0.01 and df = 129) across all the gene’s ratios. Genes that were deemed significant are reported as changed, either deletions or amplifications. As an additional method, we used samtools to create mpileup files (settings -B –d 1000000 –q 15), and subsequently, Varscan2 copynumber and Varscan2 copyCaller with default parameters to determine copy number^18^. We used the Integrated Genome Viewer **(IGV)** to visually inspect sequence reads and variant positions.^47^

#### Digital PCR confirmation of ALK amplification

The ddPCR assay was performed as described above using probes for *ALK* and *RPP30* (BioRad, Hercules, CA) for HD753 and NA12878 as control. The *ALK* result was first normalized to that of RPP30 for each sample, and then the normalized *ALK* ratio was compared for HD753 versus NA12878 to calculate a final ratio.

## ACKNOWLEDGEMENT

We would like to thank Dr. Lincoln Nadauld for his designing the gene panel, as well as Greg Jensen and Wolfgang Daum for discussion. We would also want to thank Amy Wong, Jennifer Pecson, Girish Putcha, Julie Ballard, Anagh Vora, and Alexander MacKenzie for additional comments. This work was partly supported by a National Institutes of Health / National Cancer Institute award from the Innovative Molecular Analysis Technologies program (R33CA174575).

## CONTRIBUTIONS

AS, AV, RK, SMG, YP, DM, FG, FDLV and HPJ designed, performed, and analyzed the experiments for this study. AS, AV, FDLV and HPJ wrote the manuscript. Methods originally conceived by AS, ML, and HPJ and initial development work was performed by AS, YB, ML, JZ, JS.

